# Predicting gene expression changes upon epigenomic drug treatment

**DOI:** 10.1101/2023.07.20.549955

**Authors:** Piyush Agrawal, Vishaka Gopalan, Sridhar Hannenhalli

## Abstract

**Background:** Tumors are characterized by global changes in epigenetic changes such as DNA methylation and histone modifications that are functionally linked to tumor progression. Accordingly, several drugs targeting the epigenome have been proposed for cancer therapy, notably, histone deacetylase inhibitors (HDACi) such as *Vorinostatis* and DNA methyltransferase inhibitors (DNMTi) such as *Zebularine*. However, a fundamental challenge with such approaches is the lack of genomic specificity, i.e., the transcriptional changes at different genomic loci can be highly variable thus making it difficult to predict the consequences on the global transcriptome and drug response. For instance, treatment with DNMTi may upregulate the expression of not only a tumor suppressor but also an oncogene leading to unintended adverse effect.

**Methods:** Given the pre-treatment transcriptome and epigenomic profile of a sample, we assessed the extent of predictability of locus-specific changes in gene expression upon treatment with HDACi using machine learning.

**Results:** We found that in two cell lines (HCT116 treated with Largazole at 8 doses and RH4 treated with Entinostat at 1µM) where the appropriate data (pre-treatment transcriptome and epigenome as well as post-treatment transcriptome) is available, our model distinguished the post-treatment up versus downregulated genes with high accuracy (up to ROC of 0.89). Furthermore, a model trained on one cell line is applicable to another cell line suggesting generalizability of the model.

**Conclusions:** Here we present a first assessment of the predictability of genome-wide transcriptomic changes upon treatment with HDACi. Lack of appropriate omics data from clinical trials of epigenetic drugs currently hampers the assessment of applicability of our approach in clinical setting.

## Introduction

Phenotypic state of a cell or tissue, either in normal homeostasis or in disease such as cancer, is intimately linked to its transcriptional state, which in turn is profoundly determined by its global epigenome (1). Cancer genomes display a substantially altered epigenome relative to their non-malignant counterparts. For instance, global DNA hypomethylation and focal hypermethylation, notably, at tumor suppressor gene promoters, has been noted as a general feature of many cancers (2). Accordingly, drugs that alter the epigenome have emerged as potential candidates for cancer therapy (3). Two of the most common classes of epigenome-altering drugs are DNA methyltransferase inhibitors (DNMTi), such as *Zebularine*, and Histone deacetylase inhibitors (HADCi), such as *Vorinostatis*. While DNMTi’s are standard of care in some hematological malignancies, across most cancers, the efficacy of epigenomic drugs have been mixed.

One of the many reasons adversely affecting the success of epigenomic drugs is its lack of locus specificity. Note that the intent of DNMTi, for instance, is partly to reactivate aberrantly silenced genes by demethylating their (aberrantly methylated) promoters (3,4). However, the drug is locus-agnostic and, a priori, can activate many other loci in the genome, some of which may have toxic side effects (5) and in the worst case, have pro-tumor effects; indeed, DNMTi’s are known to activate cancer testis antigens, which are known to be pro-tumor. Currently, we lack the sufficient knowledge to predict locus-specific effects of an epigenomic drug on the gene expression, to be able to develop more rational therapies. One of the facts complicating this understanding is incompletely understood interactions between different epigenomic marks. For instance, there is broad antagonism between two key mechanisms of transcriptional suppression, namely, histone modification H3K27me3 and DNA methylation (6) which may result in redistribution of one upon perturbation of the other. Because of the complex interactions between epigenomic marks as well as feed-forward and feed-back loops between the epigenome and the transcriptome, the ultimate effect of epigenomic perturbation on the global transcriptome may not be easily predictable, especially based only on the local genomic context.

Motivated by these challenges, our goal here is to assess the scope and extent to which one can predict locus-specific changes in gene expression upon treatment with an epigenomic drug such as HDACi or DNMTi, given the transcriptional and epigenomic state of the tumor sample pre-treatment. This could substantially help assess the clinical efficacy of an epigenomic drug.

## Methods

### Processing of RH4 and HCT116 sequencing data

FASTQ files were downloaded from SRA (HCT116 dataset accession: SRP113250, RH4 accession: SRP151465). We uniformly re-processed both RNA-seq and H3K27ac ChIP-seq data to minimize biases. We ran the fastqc toolkit (v0.11.9) to ensure quality. Trimgalore (v0.6.7) was run with default options to trim any adaptor sequence contamination in reads. For ChIP-seq data, bwa-mem2 (v2.2.1) (7) was used to align trimmed reads while salmon (v1.7.0) (8) was run to align trimmed reads with the –validateMappings option enabled. For ChIP-seq data, the read counts in each genomic bin (defined below) were normalized to TPM (transcripts per million) scale with genomic bin counts quantified using the Rsubread package (9). Since the RH4 ChIP-seq data contained spike-in reads from the Drosophila melanogaster genome, bwa-mem2 was used to align reads using a joint BWA index of the hg38 and dm6 genome. Thus, for the RH4 data, in addition to the library size normalization that is applied to each sample, we additionally divided the TPM values by the total number of reads that aligned to the dm6 genome assembly following the recommendation in the source publication describing the Drosophila melanogaster spike-in protocol for ChIP-seq data (10). All pseudo-aligned RNA-seq data from salmon was normalized to a TPM scale using the tximport function (v1.28.0).

### Distribution of histone marks in the genic region

We identified 1000 most upregulated and downregulated genes post drug treatment for HCT116 and RH4 cell line. The genes were selected based on Log10 Fold change i.e. Log10 Treated – Log10 Untreated. For every gene, we created 21 genomic bins to analyze the pattern of histone marks. The genomic bins include promoter region, transcription Start Site (TSS), and Gene Body (GB) region. TSS coordinates were obtained from the ENSEMBL Genes v101 database (11). We defined promoter as the 2kb region upstream to the TSS which was further divided into 10 equal-sized bins where the TSS was the single nucleotide position. Finally, the gene body was defined as the entire transcribed region and was also divided into 10 equal-sized bins. Overall, this resulted in a total of 21 bins for every gene. H3K27Ac read density was calculated in each of these 21 bins and was used to compare the up and down genes and as features for the prediction of up and down-regulated genes.

### Machine learning model to predict post-treatment transcriptional effect

We used the histone mark distribution in 21 genic bins as features to develop machine learning models to distinguish up versus downregulated genes after HDACi-treatment separately for both HCT116 and RH4 cell lines. Using the conventional 5-fold cross validation we computed Area Under Curve (AUC) as performance measure. We used a python-based library known as Scikit-learn (12) and implemented 3 different machine learning techniques which include Support Vector Machine (SVM), Random Forest (RF), and Gradient Boosting. Models were developed in 4 different categories (i) using 10 Promoter features; (ii) using single TSS feature; (iii) using 10 GB features, and lastly (iv) using all 21 features. We further performed cross cell line prediction where a model trained on one cell line data was used to predict other cell line data.

### Gene Ontology Analysis

We used clusterProfiler 4.0 (13) to identify biological processes associated with the identified up and downregulated genes. We used the following command to get the enriched significant processes: “**ego <-enrichGO(de$Entrezid, OrgDb = “org.Hs.eg.db”, ont=“BP”, readable = TRUE, minGSSize = 10, maxGSSize = 500, keyType=“SYMBOL”)**”

As there are many redundant processes, we further obtained the parent processes using the following command:

**“ego2 <-simplify(ego, cutoff=0.8, by=“p.adjust”, select_fun=min, measure = “Wang”)”**

Dotplot of the above obtained processes were created using ggplot2 library in R (14).

## Results

### Distinct patterns of epigenomic marks in gene locus between up and down regulated genes upon HDACi treatment

For each of the two cell lines (HCT116 and RH4), for the respective dosage of HDACi drugs (8 doses of Largazole for HCT116, 1 dose of 1µM Entinostat for RH4), we first identified the top 1000 up-regulated and 1000 down-regulated genes (Methods). Genes classified as up, down, and unchanged post-treatment for various doses in HCT116 and RH4 cell lines are provided in the **Supplementary Table S1 and S2 respectively**. TPM value of each gene, untreated as well as treated for various concentrations in HCT116 cell line and single concentration for RH4 is also provided in the **Supplementary Table S3 & S4** respectively.

We first identified enriched GO terms in each set of up and down-regulated genes (3 pairs of gene sets for 3 representative doses in HCT116 [4.68nM, 75nM and 300nM] and 1 pair for RH4). In general, the upregulated genes in both the cell lines were broadly enriched for the developmental and signaling processes **[Figure 1]**. The developmental process is in the direction of Epithelial to Mesenchymal Transition (EMT). In the case of HCT116 cell line, we additionally observed response to hypoxia. Likewise, processes associated with downregulated genes are broadly associated with the cell cycle and cell division whereas for RH4 cell line additional processes such as histone modification, RNA splicing were also seen **[Figure 2]**. Complete list of processes associated with the upregulated and downregulated genes in HCT116 [4.68nM, 75nM and 300nM] and RH4 cell lines are provided in the **Supplementary Table S5-S7 and S8 respectively**.

**Figure 1:**
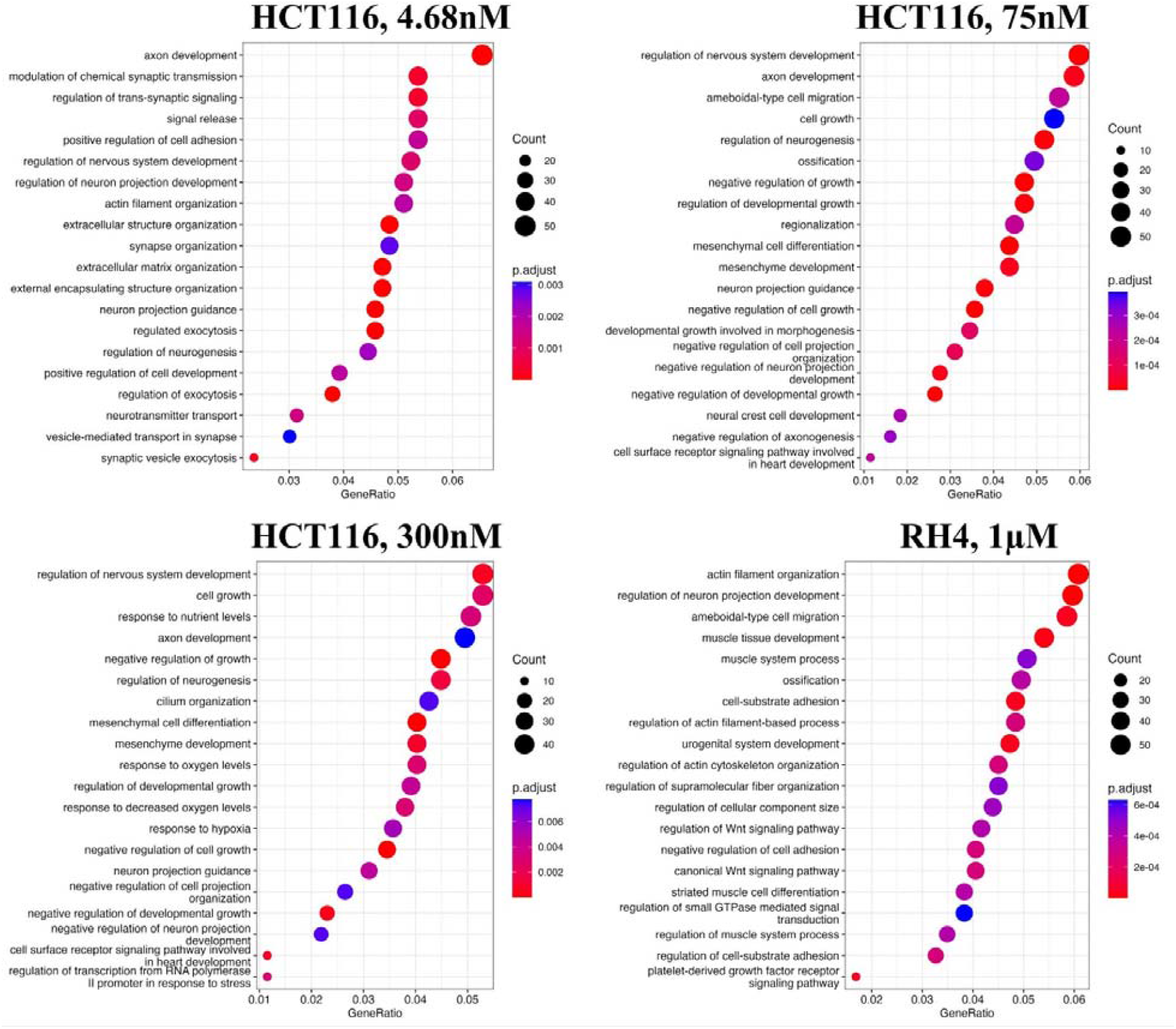
Upregulated genes are broadly enriched for the developmental and signaling processes. Top20 enriched biological processes associated with upregulated genes in HCT116 cell line after treating with epigenetic drug Largazole at 4.68nM (A); 75nM (B); 300nM (C); and Top20 enriched biological processes associated with upregulated genes in RH4 cell line after treating with epigenetic drug Entinostat at 1µM (D).

**Figure 2:**
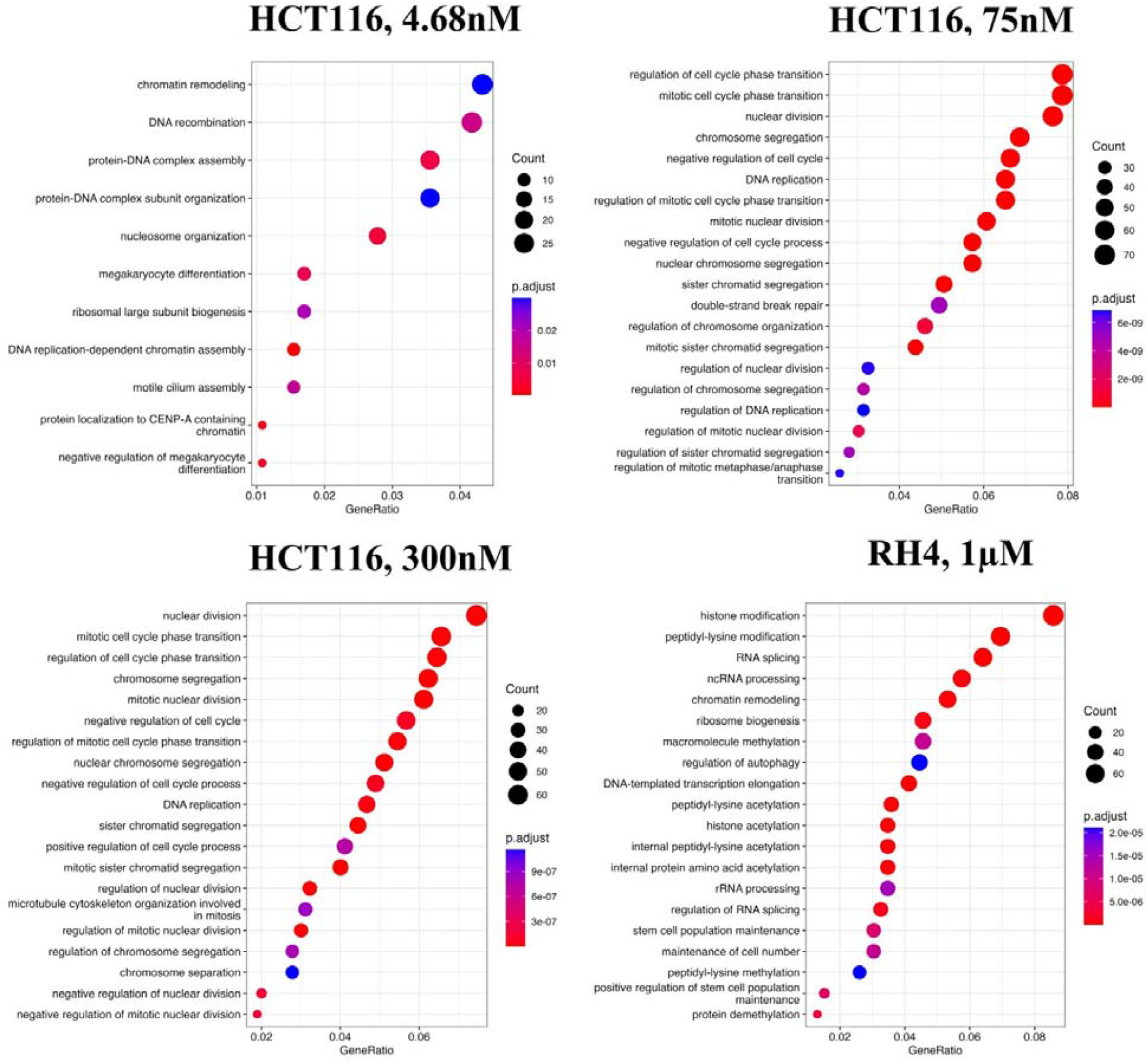
Downregulated genes are broadly enriched for the cell cycle processes. Top20 enriched biological processes associated with downregulated genes in HCT116 cell line after treating with epigenetic drug Largazole at 4.68nM (A); 75nM (B); 300nM (C); and Top20 enriched biological processes associated with upregulated genes in RH4 cell line after treating with epigenetic drug Entinostat at 1µM (D).

Next, we compared the pre-treatment epigenomic profiles of the up and down genes by plotting the H3K27Ac mark intensity (normalized read counts) in the pre-treatment sample along 21 genic bins (Methods). Distributions for three representative doses for HCT116 (4.68nM (lowest), 75nM, and 300nM (highest)) and 1µM dose for RH4 are included in **Figure 3**; all other distributions for the HCT116 cell line are provided in **Supplementary Figure S1-S5**. Overall, the following general trends emerge: (1)

**Figure 3:**
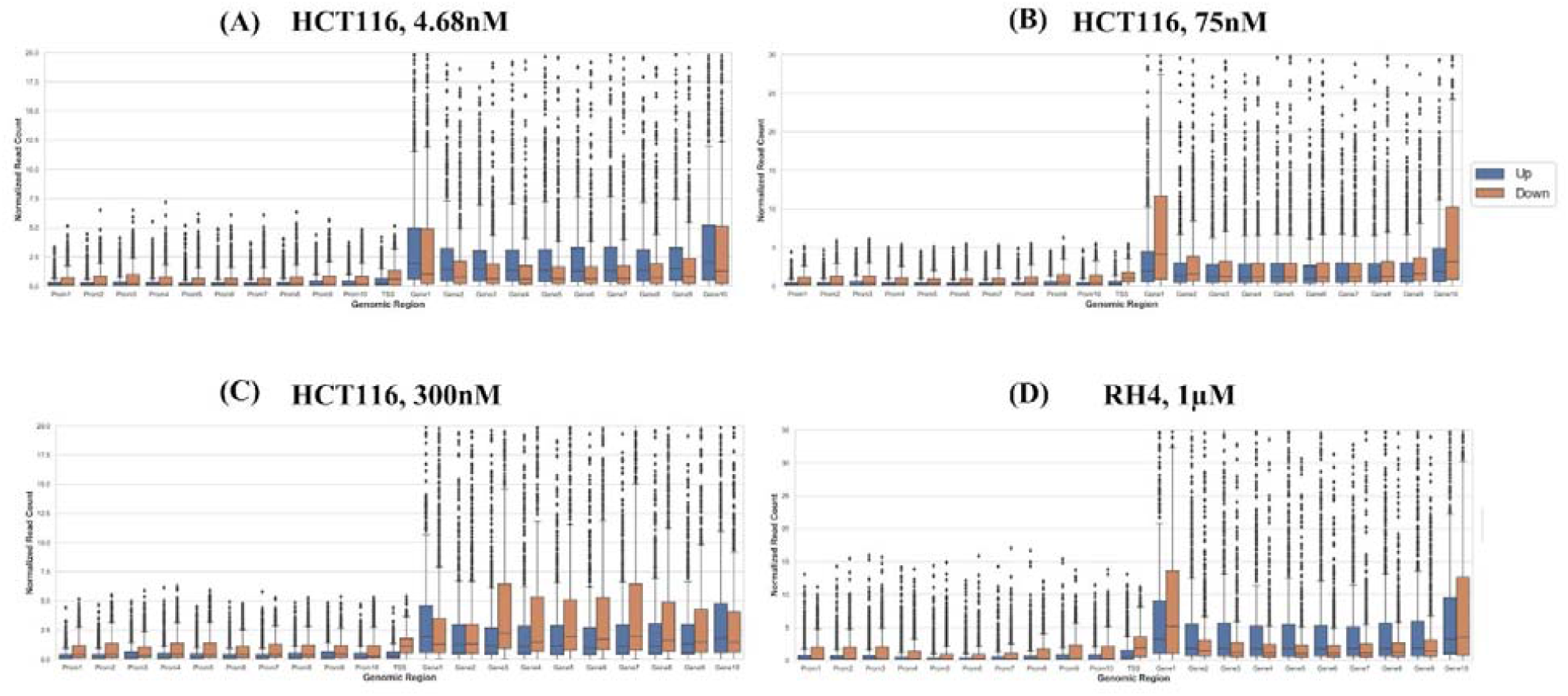
H3K27Ac mark distribution across genomic bins. Boxplot distribution of H3K27Ac marks across 21 genomic bins (10 equal sized bins of Promoter, Gene body and 1 bin of TSS) associated with upregulated (blue bars) and downregulated (brown bars) genes when HCT116 cell line is treated with Largazole at concentrations 4.68nM (A); 75nM (B); 300nM (C); and when RH4 cell line is treated with Entinostat at 1µM (D).

There is substantial variability across the bins around the genic locus in the upregulated vs downregulated H3K27Ac mark density, (2) In the upstream regions downregulated genes have a higher H3K27Ac pre-treatment, and (3) this trend is also true in gene body but only at mid and higher dosage, while (4) at low dosage the trend is opposite in gene body where the downregulated genes have lower H3K27Ac, (5) RH4 trends at 1µm dose of Entinostat most resembles the patterns at 75nM dose of Largazole in HCT116. Overall, while there is a variable pre-treatment epigenomic pattern within the gene body across cell lines, drug, and dosages, there is nevertheless sufficient differences between up and downregulated genes, motivating us to develop machine learning models to predict transcription effects given the H3K27Ac pattern at a gene locus.

### Predicting HDACi treatment impact on gene expression from the epigenome

Here, we assess whether the pre-treatment epigenetic profile at a gene locus can predict whether the gene will be up-regulated or down-regulated upon treatment with HDACi. Top 1000 up-regulated and 1000 down-regulated genes were compiled. For every gene, pre-treatment H3K27Ac read count in 21 regions relative to the gene (Methods) were used as features and three machine learning models --Support Vector Machine (SVM), Random Forest (RF), and Gradient Boosting (GB), were benchmarked based on 5-fold cross-validation and accuracy was quantified as area under the ROC curve (AUC). A separate model was benchmarked for each of the 8 drug dosages in HCT116 data. As shown in Table 1, overall, various machine learning approaches performed comparably and using all features was preferable; specifically best performance was achieved by SVM for 75nM dosage with AUC of 0.89 **(Figure 4)**. Analogous benchmarking for RH4 cell line data available at the single dosage using all features yielded comparable AUC ranging from 0.74-0.76 for the three machine learning methods. Overall, H3K27Ac signal near the gene is informative of the gene expression changes upon treatment with HDACi.

**Table 1.** Performance of various machine learning models on HCT116 and RH4 cell line testing dataset. Here the concentration of drugs is in nM and µM. P stands for Promoter; TSS stands for Transcription Start Site; GB stands for Gene Body; and All is the combination of all three features.

|  | Support Vector Machine |  |  |  | Random Forest |  |  |  | Gradient Boosting |  |  |  |
| --- | --- | --- | --- | --- | --- | --- | --- | --- | --- | --- | --- | --- |
| Con. | P | TSS | GB | All | P | TSS | GB | All | P | TSS | GB | All |
| HCT116 (4.68nM) | 0.57 | 0.61 | 0.68 | 0.71 | 0.61 | 0.68 | 0.69 | 0.74 | 0.62 | 0.69 | 0.65 | 0.73 |
| HCT116 (9.37nM) | 0.74 | 0.73 | 0.77 | 0.78 | 0.74 | 0.76 | 0.82 | 0.80 | 0.72 | 0.77 | 0.75 | 0.79 |
| HCT116 (18.75nM) | 0.60 | 0.74 | 0.74 | 0.83 | 0.68 | 0.74 | 0.74 | 0.85 | 0.66 | 0.76 | 0.74 | 0.80 |
| HCT116 (30nM) | 0.74 | 0.77 | 0.82 | 0.86 | 0.75 | 0.76 | 0.82 | 0.84 | 0.74 | 0.78 | 0.73 | 0.80 |
| HCT116 (37.5nM) | 0.82 | 0.80 | 0.85 | 0.87 | 0.83 | 0.74 | 0.85 | 0.85 | 0.82 | 0.79 | 0.78 | 0.80 |
| HCT116 (75nM) | 0.74 | 0.78 | 0.84 | 0.89 | 0.72 | 0.75 | 0.83 | 0.87 | 0.72 | 0.77 | 0.78 | 0.81 |
| HCT116 (150nM) | 0.73 | 0.79 | 0.86 | 0.87 | 0.72 | 0.76 | 0.85 | 0.87 | 0.73 | 0.79 | 0.77 | 0.82 |
| HCT116 (300nM) | 0.75 | 0.75 | 0.86 | 0.88 | 0.76 | 0.74 | 0.86 | 0.86 | 0.76 | 0.75 | 0.76 | 0.80 |
| RH4 (1μM) | 0.74 | 0.67 | 0.71 | 0.75 | 0.71 | 0.63 | 0.73 | 0.76 | 0.70 | 0.66 | 0.70 | 0.74 |

**Figure 4:**
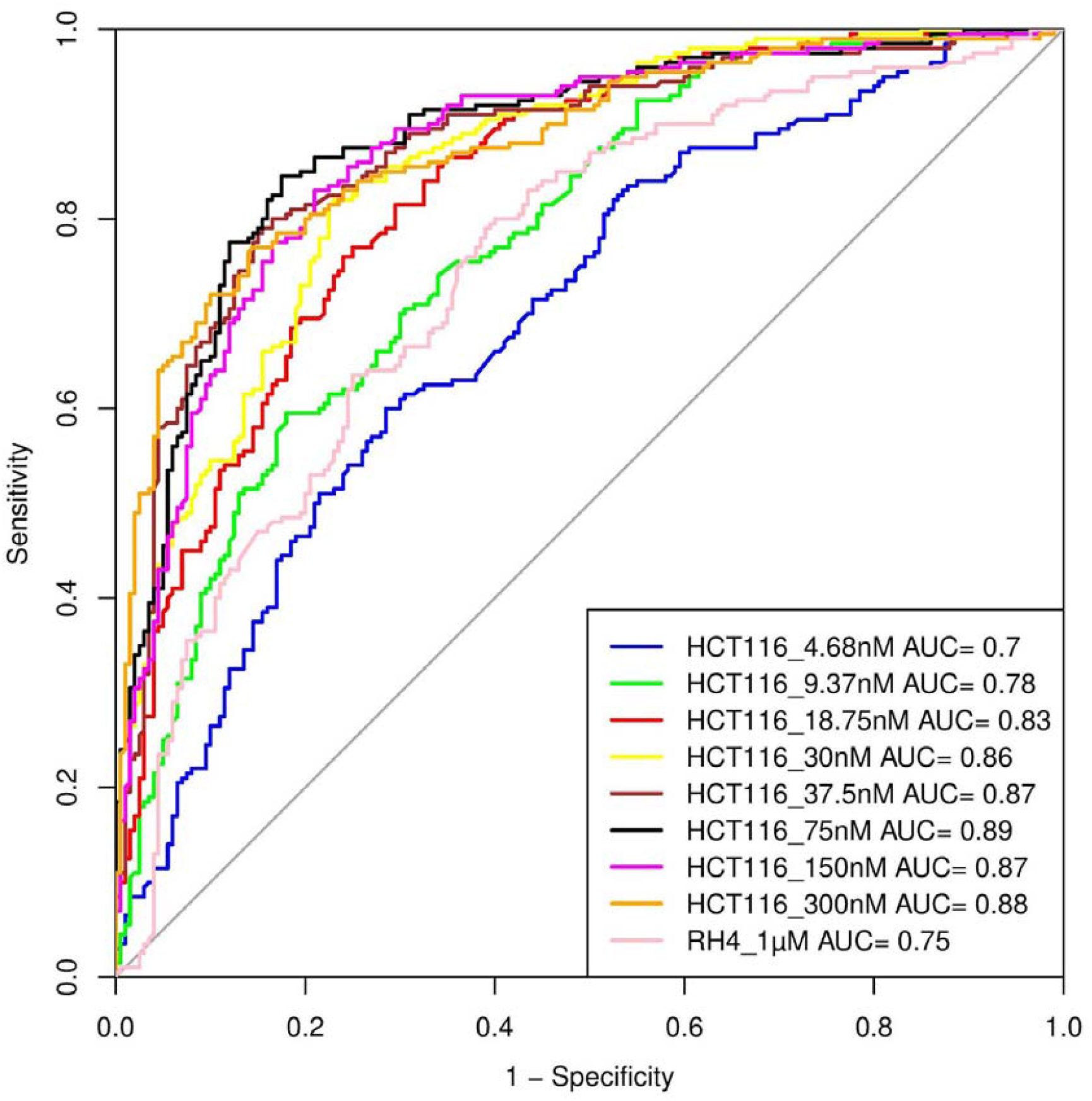
Performance of SVM model. Performance of various SVM based models in terms of Area Under Curve (AUC) at different concentrations when HCT116 cell line was treated with 8 different Largazole concentration and RH4 cell line was treated with Entinostat.

### Prediction model is generalizable across cell lines

Next, we assessed whether a model trained on one cell line to predict the transcriptional effect of a certain epigenomic drug can predict the effect in a different cell line treated with a different drug, albeit also HDACi. Toward this, first, a SVM (75nM) model trained on HCT116 cell line data was able to achieve an AUC value of 0.71 when applied to RH4 cell line data **(Figure 5A)**. Likewise, the model trained on RH4 cell line data when applied to HCT116 data achieved an AUC of 0.81 **(Figure 5B)**, supporting the cross-context generalizability of the model, consistent with similarity of epigenomic profile trends between the two cell lines as shown above **(Figure 3B & 3D)**.

**Figure 5:**
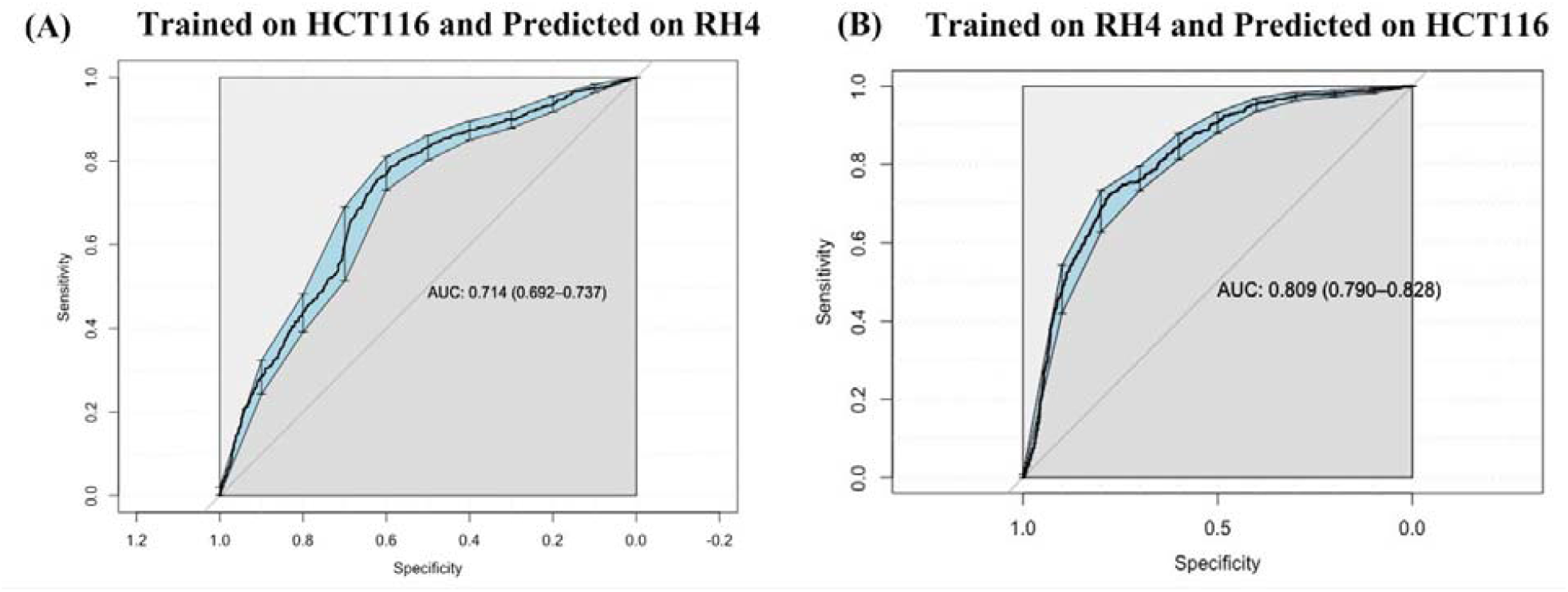
Cross cell line Prediction: (A) Performance of HCT116 data trained SVM model on RH4 cell line used as testing dataset. (B) Performance of RH4 data trained SVM model on HCT116 cell line used as testing dataset.

### Cell line-specific expression changes are reflected in their epigenome

Next, we specifically assessed whether the context-specific differences across the two cell lines in their HDACi-induced gene expression are reflected in their context-specific pre-treatment epigenomic profile in the gene locus. Toward this we compared the data for HCT116 treated with 75nM Largazole with RH4 treated with 1µM of Entinostat. For each cell line we applied stringent criteria to identify genes which were upregulated in one cell line and downregulated in another cell line. We selected those genes whose fold change > 3 in one cell line and < 1/3 in another. This resulted in two gene sets: (1) 73 genes upregulated in HCT116 and downregulated in RH4, and (2) 184 genes upregulated in RH4 and downregulated in HCT116. To normalize for cell line-specific differences in H3K27Ac, we z-scored the cross-bin H3K27Ac signal for each gene. Then for these two gene sets, we plotted the normalized H3K27Ac intensities along the 21 genic bins, comparing two cell lines features. As shown below, with few exceptions, for the first gene set, HCT116 genes show higher concentration of H3K27Ac marks and for the second gene set, the opposite is true **(Figure 6B)**, consistent with the patterns in **Figure 3**.

**Figure 6:**
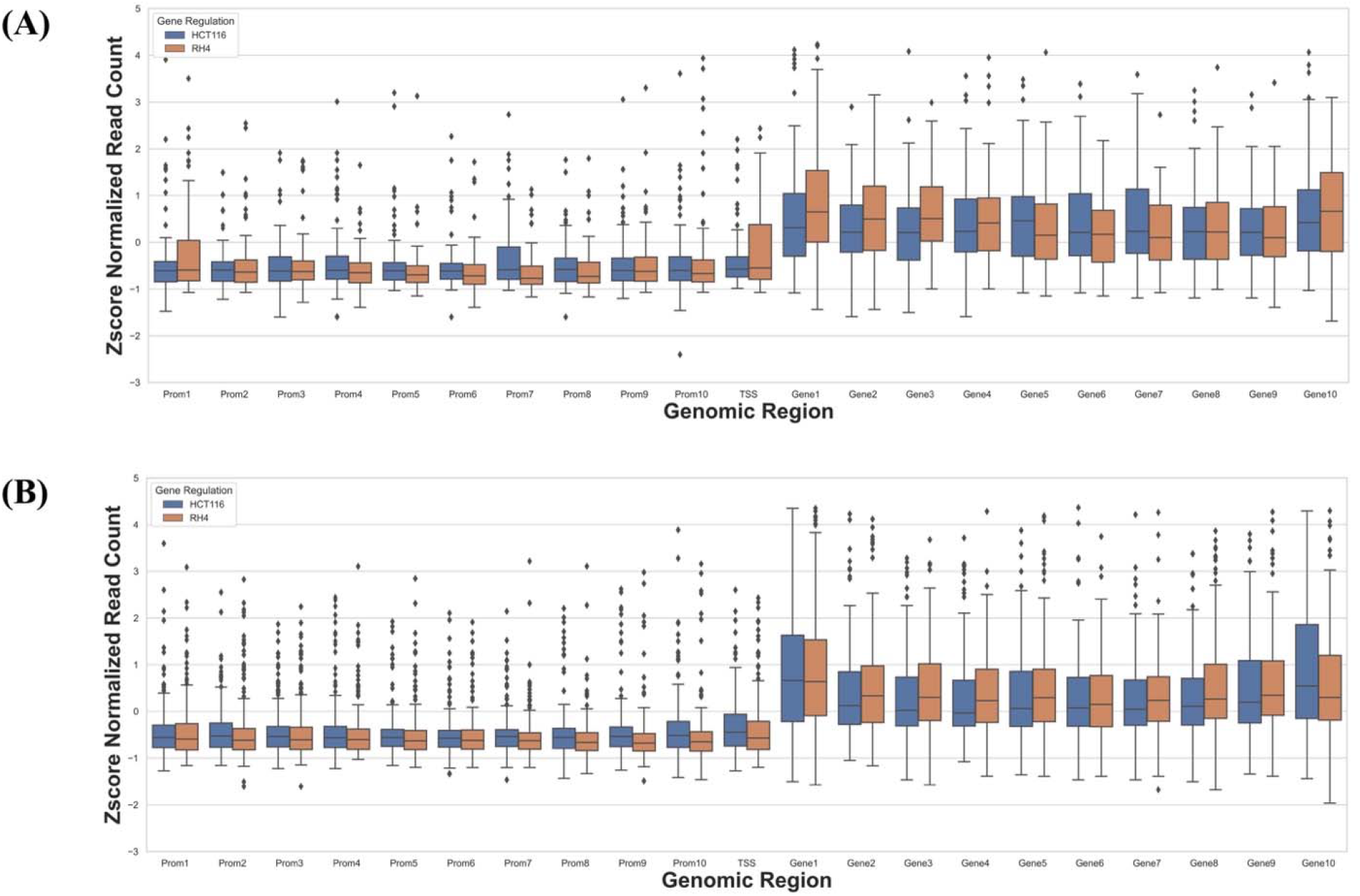
H3K27Ac mark distribution across genomic bins in two cell lines. Boxplot distribution of H3K27Ac marks across 21 genomic bins (10 equal sized bins of Promoter, Gene body and 1 bin of TSS) associated with genes with positive log fold change in HCT116 but negative log fold change in RH4 cell line (A) and Genes with positive log fold change in RH4 but negative log fold change in HCT116 cell line (B).

## Discussion

Epigenetic dysregulation is a key characteristic of cancers. Number of mutations have been observed in the genes encoding epigenetic modifiers such as DNA methylation and histone modification enzymes (15). Accordingly, efforts have been made in targeting epigenetic regulators (16). At present, seven epigenetics-targeting drugs have been approved by the FDA (17). However, there are certain challenges associated with this class of drugs, limiting their success. Some of the key challenges include (i) Different epigenetic mutations are associated with different cancer types; (ii) The same gene may have opposite function in tumorigenesis of different cancers. For example, *EZH2* deficiency causes myeloid malignancies (18) whereas gain-of-function causes B cell lymphomas (19); (iii) Another major issue is the selectivity of these drugs. For example, 30 enzymes of the KDM family with similar JMJC domain belong to 5 subfamilies. These enzymes demethylate different histone residues. Hence, drugs targeting these are broad spectrum affecting multiple KDM subfamilies and histone marks with potentially unintended consequences (20); (iv) Yet another issue with epigenetics-targeting drugs, focused in this work, is the selectivity of genomic loci. For instance, a HDACi can both increase as well as decrease histone acetylation in different genomic loci and can thus upregulate certain genes while downregulating others, again with unintended consequences.

Here, we tried to address the selectivity issue by developing a machine learning model based on pretreatment histone mark. We establish in two cell lines that the locus-specific effect of HDACi treatment on gene expression can be predicted to a reasonable accuracy from the pre-treatment histone acetylation pattern at a gene locus, and the model appears to be generalizable across cell lines. While the current study is promising and may potentially be applied to personalized therapy by predicting the transcriptomic consequence of HDACi treatment, there are few limitations which need to be addressed. Our predictive model is based only on the H3K27ac mark. Several other marks such as H3K9ac, H3K4me3, H3K27me3, etc. should be incorporated in such modeling approaches in the future as and when such data become available. Our model was assessed only in cell lines and its efficacy in bulk tumor data representing the tumor microenvironment remains to be assessed. Last but not the least, pre-and post-treatment tumor epigenetic and transcriptomic data in clinical and pre-clinical models are still lacking, required for assessing clinical applicability of our approach.

## Acknowledgement

This work utilized the computational resources of the NIH HPC Biowulf cluster. Authors are also thankful to the other lab members for their valuable suggestions. This work was supported by the Intramural Research Program of the National Cancer Institute, Center for Cancer Research.

## Author contribution

VG download and processed the data. PA, and SH perform the analysis. PA and SH perform the statistical analysis. PA, VG and SH wrote the manuscript. PA and SH supervised the study. All authors read the article and approved the submitted version.

## Data and Code Availability

All the codes used in the current study are deposited at GitHub (https://github.com/hannenhalli-lab/Epigenetic_Project/). Further inquiries can be directed to the corresponding authors.

