## Supplementary Figures for "Predicting gene expression changes upon epigenomic drug treatment"

1. Sridhar Hannenhalli

Building 10, Room 2-6300

Bethesda, MD 20814

2. Piyush Agrawal

Building 10, Room 2-6300

Bethesda, MD 20814

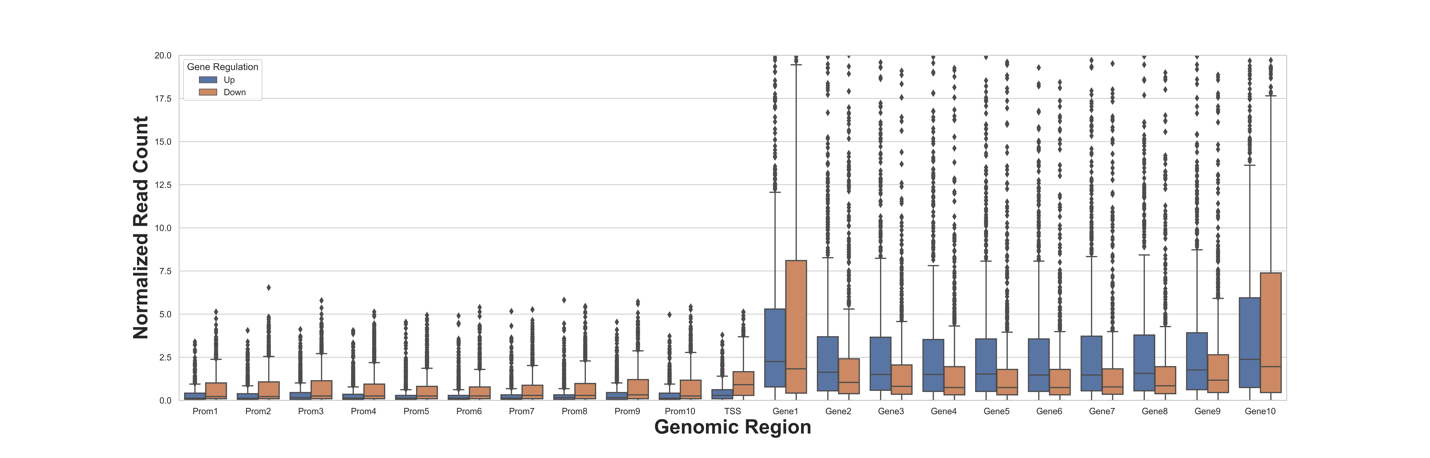

**Figure S1: H3K27Ac mark distribution across genomic bins.** Boxplot distribution of H3K27Ac marks across 21 genomic bins (10 equal sized bins of Promoter, Gene body and 1 bin of TSS) associated with upregulated (blue bars) and downregulated (brown bars) genes when HCT116 cell line is treated with Largazole at concentrations 9.37nM.

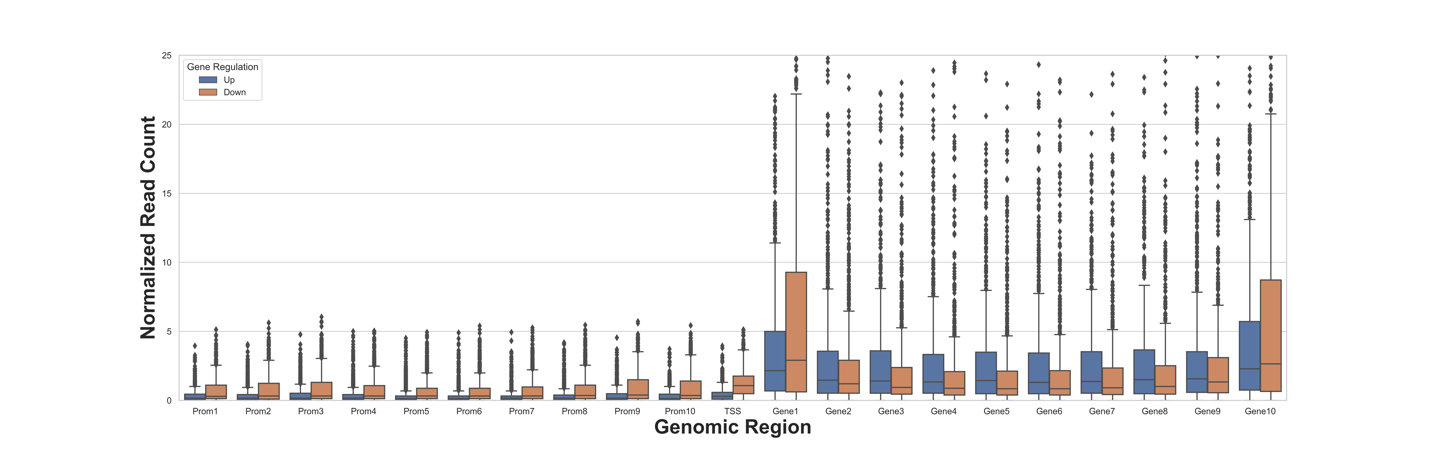

**Figure S2: H3K27Ac mark distribution across genomic bins.** Boxplot distribution of H3K27Ac marks across 21 genomic bins (10 equal sized bins of Promoter, Gene body and 1 bin of TSS) associated with upregulated (blue bars) and downregulated (brown bars) genes when HCT116 cell line is treated with Largazole at concentrations 18.75nM.

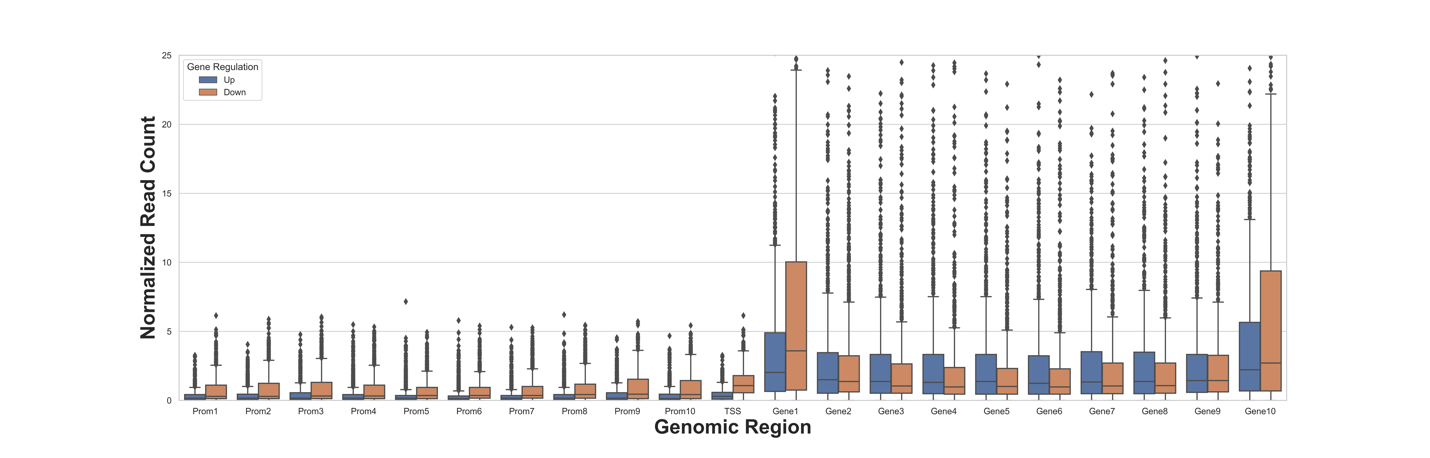

**Figure S3: H3K27Ac mark distribution across genomic bins.** Boxplot distribution of H3K27Ac marks across 21 genomic bins (10 equal sized bins of Promoter, Gene body and 1 bin of TSS) associated with upregulated (blue bars) and downregulated (brown bars) genes when HCT116 cell line is treated with Largazole at concentrations 30nM.

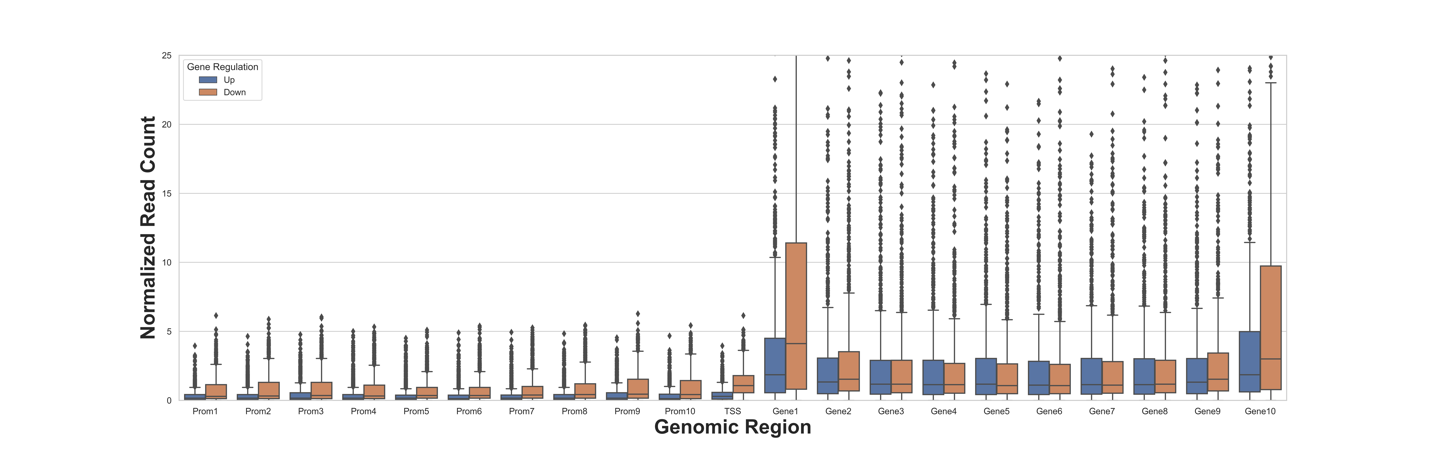

**Figure S4: H3K27Ac mark distribution across genomic bins.** Boxplot distribution of H3K27Ac marks across 21 genomic bins (10 equal sized bins of Promoter, Gene body and 1 bin of TSS) associated with upregulated (blue bars) and downregulated (brown bars) genes when HCT116 cell line is treated with Largazole at concentrations 37.50nM.

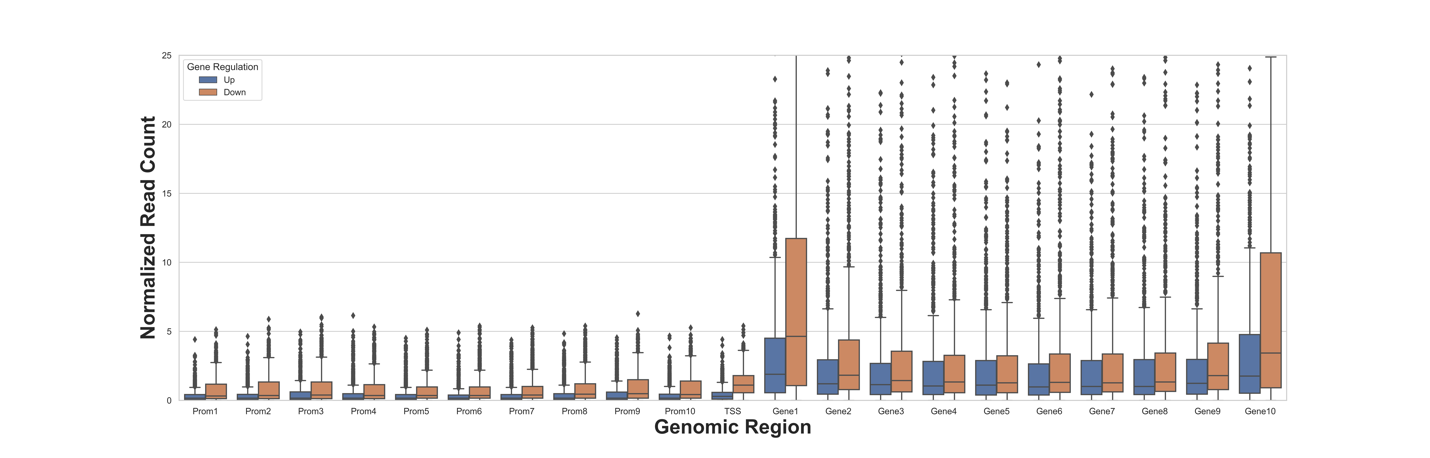

**Figure S5: H3K27Ac mark distribution across genomic bins.** Boxplot distribution of H3K27Ac marks across 21 genomic bins (10 equal sized bins of Promoter, Gene body and 1 bin of TSS) associated with upregulated (blue bars) and downregulated (brown bars) genes when HCT116 cell line is treated with Largazole at concentrations 150nM.
